# Western diet dampens T regulatory cell function to fuel hepatic inflammation in nonalcoholic fatty liver disease

**DOI:** 10.1101/2023.03.23.533977

**Authors:** Sudrishti Chaudhary, Ravi Rai, Pabitra B. Pal, Dana Tedesco, Aatur D. Singhi, Satdarshan P. Monga, Arash Grakoui, Smita S. Iyer, Reben Raeman

**Affiliations:** Division of Experimental Pathology, Department of Pathology, University of Pittsburgh, Pittsburgh, PA; Pittsburgh Liver Research Center, University of Pittsburgh Medical Center and University of Pittsburgh School of Medicine, Pittsburgh, PA USA; Division of Infectious diseases, Emory Vaccine Center, Division of Microbiology and Immunology, Emory University School of Medicine, Atlanta, Georgia, USA; Division of Anatomic Pathology, Department of Pathology, University of Pittsburgh School of Medicine, Pittsburgh, PA USA; Division of Gastroenterology, Hepatology and Nutrition, Department of Medicine, University of Pittsburgh School of Medicine, Pittsburgh, PA USA; Emory National Primate Research Center, Atlanta, Georgia, USA

**Author notes:** Corresponding Author: Reben Raeman, M.S., Ph.D. Assistant Professor, 200 Lothrop Street, S408 BST, Pittsburgh, PA 15261. Contributed equally (Sudrishti Chaudhary, Ravi Rai, Pabitra B. Pal). ***Conflict of interest:*** The author declares no conflicts of interest. **Author Contributions.** Conceptualization: RR; Methodology: RR, SC, RPR, PBP, ADS; Investigation: RPR, SC, PBP, DT, RR, SSI; Visualization: RPR, SC, PBP, RR, SSI; Funding acquisition: RR; Project administration: RR, RPR, SC, PBP; Supervision: RR; Writing – original draft: SC, RR; Writing – review & editing: RR, SC, SSI, AG, DT, SPM, ADS.

**Keywords:** NAFLD, NASH, Fibrosis, Inflammation, T regulatory cells

## Abstract

**Background and aims:** The immunosuppressive T regulatory cells (Tregs) regulate immune responses and maintain immune homeostasis, yet their functions in nonalcoholic fatty liver disease (NAFLD) pathogenesis remains controversial.

**Methods:** Mice were fed a normal diet (ND) or a western diet (WD) for 16 weeks to induce NAFLD. Diphtheria toxin injection to deplete Tregs in Foxp3^DTR^ mice or Treg induction therapy in WT mice to augment Treg numbers was initiated at twelve and eight weeks, respectively. Liver tissues from mice and NASH human subjects were analyzed by histology, confocal imaging, and qRT-PCR.

**Results:** WD triggered accumulation of adaptive immune cells, including Tregs and effector T cells, within the liver parenchyma. This pattern was also observed in NASH patients, where an increase in intrahepatic Tregs was noted. In the absence of adaptive immune cells in Rag1 KO mice, WD promoted accumulation of intrahepatic neutrophils and macrophages and exacerbated hepatic inflammation and fibrosis. Similarly, targeted Treg depletion exacerbated WD-induced hepatic inflammation and fibrosis. In Treg-depleted mice, hepatic injury was associated with increased accumulation of neutrophils, macrophages, and activated T cells within the liver. Conversely, induction of Tregs using recombinant IL2/αIL2 mAb cocktail reduced hepatic steatosis, inflammation, and fibrosis in WD-fed mice. Analysis of intrahepatic Tregs from WD-fed mice revealed a phenotypic signature of impaired Treg function in NAFLD. *Ex vivo* functional studies showed that glucose and palmitate, but not fructose, impaired the immunosuppressive ability of Treg cells.

**Conclusions:** Our findings indicate that the liver microenvironment in NAFLD impairs ability of Tregs to suppress effector immune cell activation, thus perpetuating chronic inflammation and driving NAFLD progression. These data suggest that targeted approaches aimed at restoring Treg function may represent a potential therapeutic strategy for treating NAFLD.

**Lay summary:** In this study, we elucidate the mechanisms contributing to the perpetuation of chronic hepatic inflammation in nonalcoholic fatty liver disease (NAFLD). We show that dietary sugar and fatty acids promote chronic hepatic inflammation in NAFLD by impairing immunosuppressive function of regulatory T cells. Finally, our preclinical data suggest that targeted approaches aimed at restoring T regulatory cell function have the potential to treat NAFLD.

## INTRODUCTION

Nonalcoholic fatty liver disease (NAFLD) is a progressive form of inflammatory liver disease encompassing a spectrum of liver pathology ranging from simple steatosis (NAFL, nonalcoholic fatty liver) to steatohepatitis and fibrosis (NASH, nonalcoholic steatohepatitis).^1^ NAFL affects one-third of the Western population, and nearly 20% people with NAFL go on to develop NASH.^1^ People with NASH are at a higher risk for developing cirrhosis and ultimately hepatocellular carcinoma (HCC). NASH-fibrosis is the second leading indication and most rapidly increasing indication for liver transplantation in the United States.^2, 3^ Despite the compelling evidence for immune dysfunction in the development and progression of NAFL to NASH and cirrhosis, the mechanisms driving immune activation and the cellular players contributing to the resulting inflammatory immune response are still unclear.

Recent evidence implicates T cells in promoting hepatic inflammation in NAFLD, yet the role played by various T cell subsets are not well understood.^4, 5^ T cells consist of inflammatory effector and immunosuppressive cell subsets, both with multifaceted roles in tissue injury and repair. In chronic tissue injury, effector T cells produce inflammatory cytokines, chemokines, and cytolytic molecules that perpetuate inflammation and injury. Conversely, immune regulatory T cells suppress these inflammatory responses, promoting tissue repair and limiting tissue injury. The interplay between these T cell subsets ultimately determines the extent of tissue injury in chronic inflammatory diseases, such as NAFLD.^6–9^

T regulatory cells (Tregs), a specialized subset of CD4 T cells expressing the transcription factor Foxp3, are crucial for maintaining immune tolerance, preventing autoimmunity, and limiting inflammatory responses in chronic inflammatory diseases. Tregs utilize several mechanisms, such as inhibiting effector T cell proliferation via IL-2 deprivation, secreting inhibitory cytokines like IL-10, IL-35, and TGFβ, and producing adenosine, to suppress T cell activation. Additionally, Tregs use CTLA-4 to block antigen-presenting cell function and granzyme and/or perforin for effector T cell cytolysis. Therefore, loss or functional impairment of Tregs is associated with several immunological and chronic inflammatory disease.^6, 10^

Despite their importance in regulating tissue immune homeostasis and suppressing inflammation in chronic inflammatory diseases, the role of Tregs in NAFLD remains unresolved. Clinical studies in NAFLD patients have yielded conflicting results linking both enrichment and decrease in Tregs in the liver and peripheral blood with disease severity.^11–16^ In mouse models where high fat, high fructose, and high cholesterol (Western diet, WD) diet was used to induce NAFLD, most studies reported enrichment of effector T cells as well as Tregs in the liver.^12, 17–19^ Increase in Treg numbers in the liver was also reported in NASH-HCC models where choline deficient high fat diet along with chemical inducers of HCC were used to induce liver injury.^20^ These findings were contradicted by a recent study reporting a proinflammatory role of Tregs in NAFLD by reporting that adoptive transfer of CD4+CD25+ Tregs exacerbate WD-induced liver injury.^21^ Since CD25 is also expressed by activated T cells and is not a specific marker for Tregs,^22^ these findings underscore the need for definitive studies to understand the role of Tregs in NAFLD progression.

Our study provides evidence supporting the anti-inflammatory role of Tregs in NAFLD pathogenesis. Our in vivo and ex vivo studies revealed that glucose and palmitic acid impair the immunosuppressive function of hepatic Tregs, leading to partial suppression of effector immune cells and perpetuation of hepatic inflammation. However, compensatory recruitment and accumulation of Tregs in the liver can prevent rapid progression of NAFLD despite their functional impairment. Collectively, our work uncovers a novel mechanism by which dietary components prevent immune regulation by Tregs, allowing chronic inflammation to persist in NAFLD.

## METHODS

### Mice

Adult male C57BL/6J and *Rag1^-/-^* mice were obtained from The Jackson Laboratory (Bar Harbor, ME). Foxp3^DTR^ knock-in mice were a gift from Dr. Rafi Ahmed.^23^ Age- and sex-matched littermates at six-weeks of age were used for all studies. The mice were maintained at the University of Pittsburgh Division of Animal Resources, and animal studies were conducted in accordance with protocols approved by the Institutional Animal Care and Use Committee at the University of Pittsburgh.

### NAFLD diet

To induce NAFLD, six-week-old male C57BL/6J, Foxp3^DTR^, and Rag1^-/-^ mice and their littermate controls were fed a western diet (WD) *ad libitum* for 16 weeks as described previously ^4, 24^. The WD (TD.130885; Harlan Laboratories) consisted of 0.2% cholesterol, 20% protein, 43% CHO, 23% fat (6.6% trans-fat), and 2.31% fructose.^24^ The normal diet (ND) comprised of 16% protein, 61% carbohydrate and 7.2% fat.

### Ex vivo analysis of Treg function

Splenic CD4 T cells from WT mice were enriched using naive untouched CD4 T cell isolation kit as per manufacturer’s instructions (Miltenyi Biotec, cat. No. 130-104-453). Isolated CD4 T cells (1 x10^6^ cells) were incubated at 37^0^ C with Dynabeads™ Mouse T-Activator CD3/CD28 for T-Cell Expansion and Activation (Thermo scientific, cat. No. 11456D,) and treated with glucose (25mM, Fisher Scientific, cat. no. D16-1), fructose (25mM, Fisher Scientific, cat. no. 161355000) or palmitic acid (400uM, Millipore Sigma, cat. no. P0500-10G) for 24-hours. Untreated CD4 T cells served as controls. Following 24 h incubation, cells were stimulated with eBioscience Cell Stimulation cocktail (eBioscience, cat. No. 00-4970-93) for 1 h, followed by incubation for an additional 4 h in the presence GolgiPlug (BD Biosciences, cat. No. 555029) and GolgiStop (BD Biosciences, cat. No. 554724). Cells were stained with a panel of fluorochrome-conjugated antibodies and viability stain (BD Biosciences, San Jose, CA) and acquired on FACS Canto II (BD) equipped with three lasers and data were analyzed by using FlowJo software as described above. Fluorochrome-conjugated antibodies against CD4 (clone RM4-5) and IL10 (clone JES5-16E3) were purchased from BD Biosciences (San Jose, CA), and Foxp3 (clone FJK-16s) and Live/Dead Aqua were purchased from Thermo Fisher Scientific (Rockford, IL).

### Treg depletion

A cohort of Foxp3^DTR^ mice fed a WD for 16 weeks were randomized to receive weekly intraperitoneal injections (IP; 50ug/Kg) of DT or saline for four weeks starting at week twelve ^25^. Diphtheria toxin was purchased from Millipore Sigma (St. Louis, MO) and was reconstituted according to the manufacturer’s protocol (Catalog: D0564).

### *In vivo* Treg induction

For the Treg induction studies, we used the established *in vivo* Treg induction cocktail consisting of recombinant murine IL2 (0.01 ug/gm, Peprotech, Catalog 212-12) complexed to anti-IL2 antibody (0.05 ug/gm; clone Jes6-1A12; BioXCell).^26^ The cocktail or PBS was administered twice weekly for four-weeks, starting at week twelve of WD-feeding.

### Human tissue

De-identified, fixed explanted liver tissue sections from disease controls (n = 10) and NASH patients (n = 10) were obtained from the Biospecimen Processing and Repository Core at the Pittsburgh Liver Research Centre.

### Statistical analysis

One-way ANOVA with post-hoc test as well as two-tailed Student’s t test were employed, as appropriate, to investigate statistical differences. A *p* value < 0.05 was considered statistically significant. Data shown are representative of 3 independent experiments. Statistical analyses were performed using Prism 9.0 (San Diego, CA, USA).

## RESULTS

### WD feeding promotes NAFLD progression in WT mice

To determine whether WD-fed C57B/6J mice (WT) is a reliable model of progressive NAFLD, we examined liver histology as well as biochemical markers of NAFLD in WT mice fed a WD for eight and sixteen weeks (**Fig. 1A-I**). Eight weeks of WD consumption significantly increased body weight as well as liver and visceral fat weight expressed as percentage of body weight (**Fig. 1E**). Histologic staining of the liver tissue revealed mild centrilobular micro and macro vesicular steatosis as well as mild lobular inflammation indicated by lobular infiltration of immune cells (**Fig. 1B**). Serum AST and ALT levels were significantly higher in the mice fed a WD for eight weeks relative to ND fed mice (**Fig. 1D**). However, eight weeks of WD feeding did not induce hepatic fibrosis as indicated by absence of marked increase in collagen deposition in the liver tissue sections stained with Sirius red (**Fig. 1C**).

**Figure 1.**
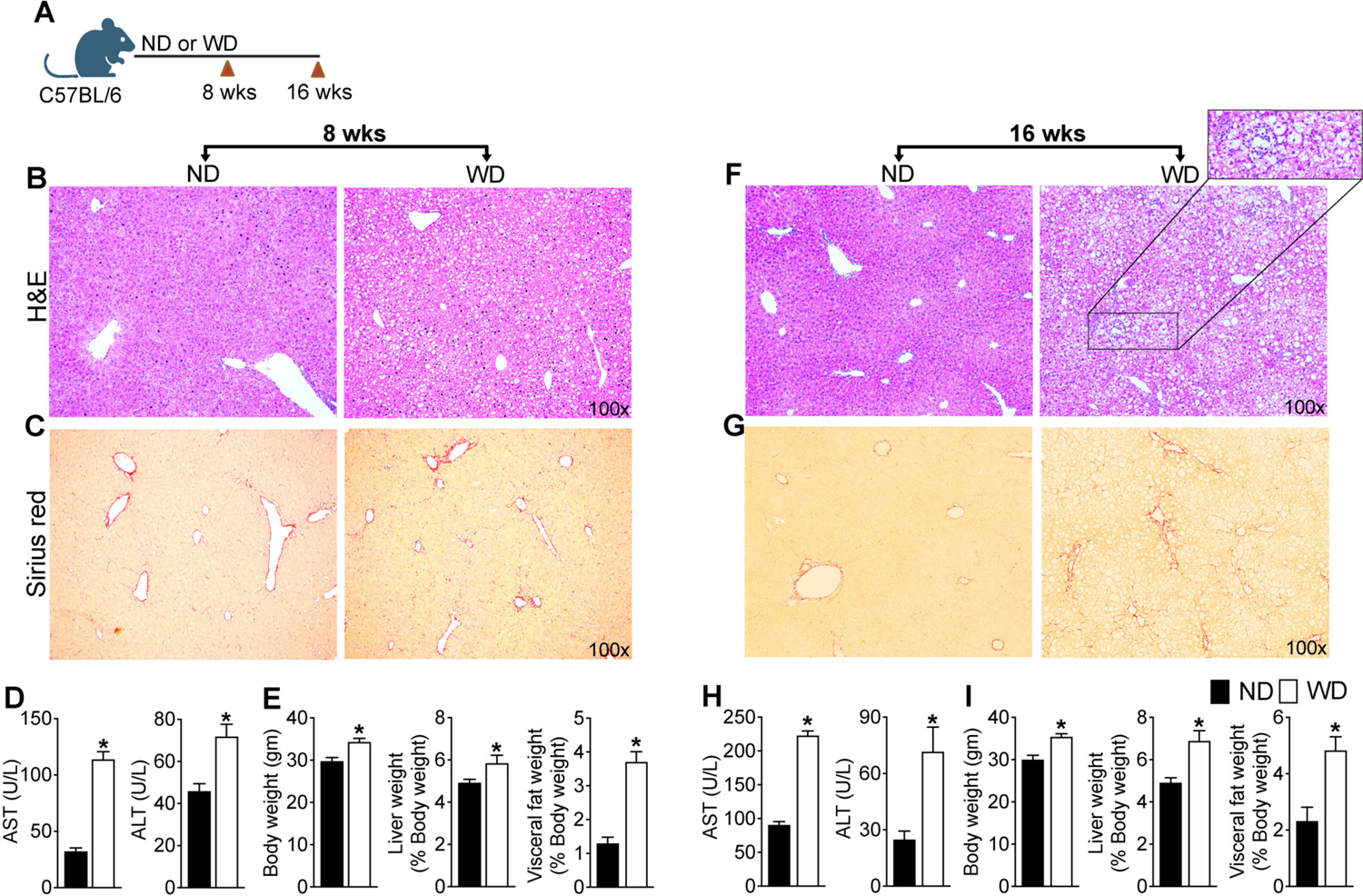
WD feeding promotes NAFLD progression in WT mice. **(A)** Schematic of study J (WT) mice were fed western diet (WD) or normal diet (ND) for 8 and 16 weeks. Representative photomicrographs of Hematoxylin and Eosin (H&E) and Sirius Red-stained liver tissue sections at **(A-B)** eight-weeks and **(E-F)** sixteen-weeks. Serum AST and ALT levels at **(C)** eight-weeks and (**G**) sixteen-weeks. Body weight and liver and visceral fat weight expressed as percentage of body weight at **(D)** eight-weeks and **(H)** sixteen-weeks. Data are representative of 3 independent experiments (n = 5 - 8 mice per group). Data are presented as mean ± SEM. Asterisks indicate significant differences (p < 0.05) between ND- and WD-fed mice.

In contrast, mice fed a WD for sixteen weeks developed severe hepatosteatosis and prominent hepatocyte ballooning and lobular inflammation demonstrating the progressive nature of the disease (**Fig. 1F**). Sixteen weeks of WD feeding enhanced hepatic fibrosis indicated by increased collagen deposition in the Sirius red stained liver tissue sections (**Fig. 1G**). Consistent with the liver pathology, sixteen weeks of WD feeding further increased serum ALT and AST levels suggesting significant disease progression (**Fig. 1H**). Disease progression in WD-fed mice correlated with increased body weight as well as liver and fat weight expressed as percentage of body weight (**Fig. 1I**). Together these data demonstrate that WD-fed WT mice is a progressive model of NAFLD and develop key histologic and biochemical features of human NASH including hepatosteatosis, inflammation, and fibrosis when fed a WD for sixteen weeks. It should be noted that NAFLD is a complex heterogeneous disease and NAFLD progression in human adults requires prolonged consumption of unhealthy diet. Thus, while WD-fed WT mice may not represent the entire spectrum of human NAFLD, the model closely recapitulates clinical characteristics of NAFLD progression in human adults.

### NAFLD progression in mice fed a WD is associated with aberrant activation of hepatic and systemic T cells

After establishing that WD-fed WT mice is a progressive model of NAFLD that closely recapitulates human disease, we next examined intrahepatic lymphocytes in mice fed the WD for sixteen weeks (**Fig. 2A**). We focused on T cell compartment as previous pre-clinical studies including our own underscore the importance of T cells in promoting NAFLD progression. ^4, 12, 17–19^ In contrast to previous reports,^21, 27^ WD-feeding did not alter the percentage of intrahepatic CD4 and CD8 T cells, but significantly increased the total numbers of CD4 and CD8 T cells in the liver of WD-fed mice relative to ND-fed controls (**Fig. 2B-D**). Further analysis revealed the enrichment of CD4 T cells expressing T cell activation markers CD44 and PD-1 in the livers of WD fed mice suggesting that WD consumption increases hepatic CD4 T cell activation (**Fig. 2E-H**). Interestingly, WD-feeding also increased the percentage and total numbers of Foxp3+ T regulatory (Tregs) cells in the liver (**Fig. 2I-J**). In contrast to the liver, we did not observe changes in either percentages or total numbers of CD4 and CD8 T cells in the spleen, suggestive of increased T cell recruitment to the liver (**Fig. 2K-L**). WD feeding, did however, enrich CD44 and PD-1 double positive CD4 T cells in the spleen of WD fed mice, which paralleled the enrichment of Foxp3+ Treg cells in the spleen (**Fig. 2M-N**). Enrichment of Foxp3+ Treg cells in liver tissue sections from individuals with non-alcoholic steatohepatitis (NASH) compared to disease controls suggests that Foxp3+ Treg cell enrichment also occurs in human NASH (**Fig. 2O**). Collectively, these data suggest that WD-feeding results in the heightened activation of hepatic T cells which parallels a compensatory increase in Foxp3+ Treg cells.

**Figure. 2.**
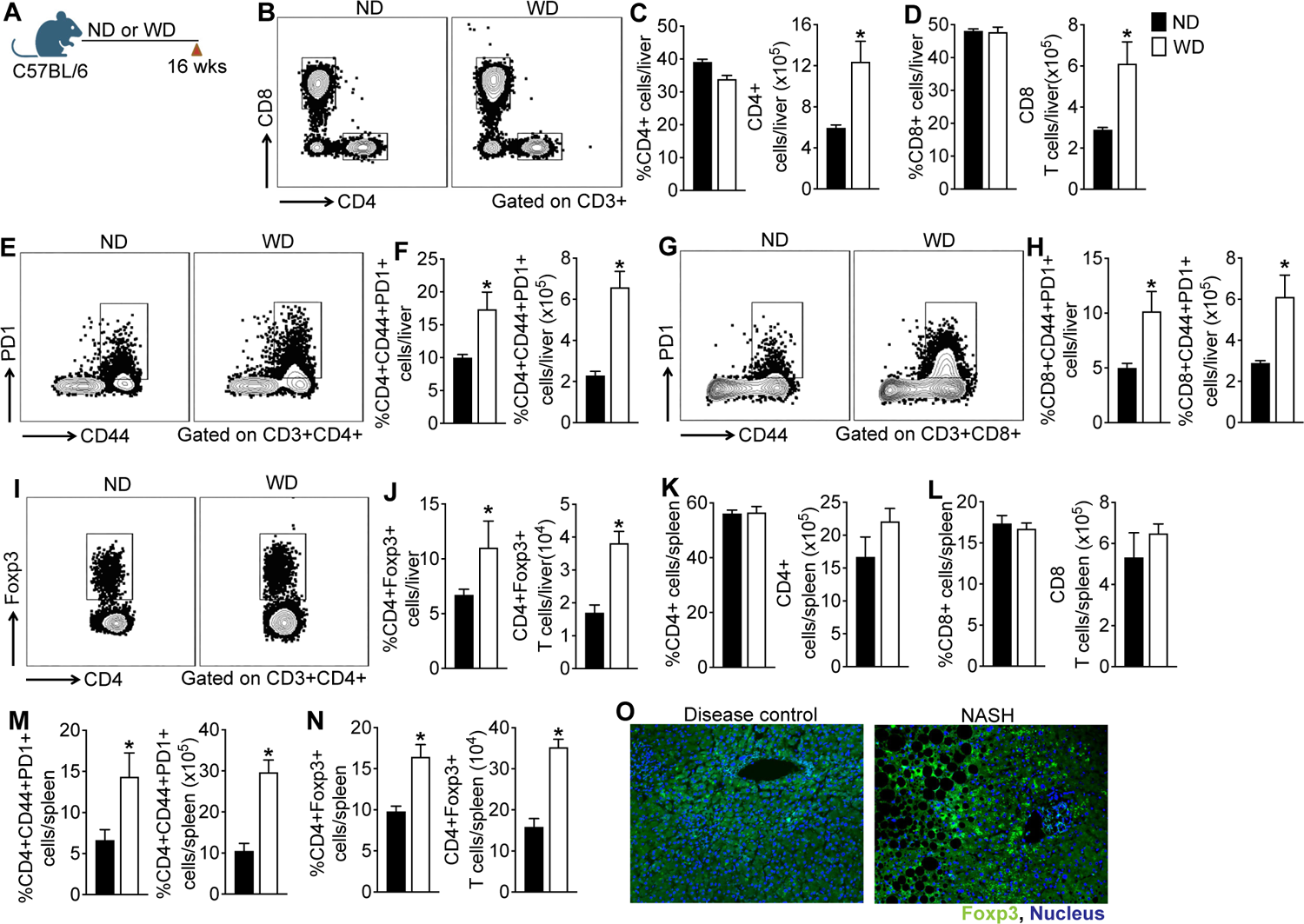
NAFLD progression in mice fed a western diet is associated with aberrant activation of hepatic and systemic T cells. (A) Schematic of study design. WT mice were fed a ND or WD for sixteen-weeks and intrahepatic lymphocytes were analyzed by FACS. **(B**) Representative flow plots show percent of CD4 T cells in the liver. Bar graphs show percent and total number of (**C**) CD4 and (**D**) CD8 T cells in the liver (n = 5 mice per group). **(E)** Representative flow plots show percent of PD-1+CD44+ CD4 T cells and **(F)** bar graphs show percent and total number of PD-1+CD44+ CD4 T cells in the liver. **(G)** Representative flow plots show percent of PD-1+CD44+ CD8 T cells and **(H)** bar graphs show percent and total numbers of PD-1+CD44+ CD8 T cells in the liver. **(I)** Representative flow plots show percent of Foxp3+ CD4 T cells and **(J)** bar graphs show percent and total number of Foxp3+ CD4 T cells in the liver. Bar graphs show percent and total number of (**K**) CD4 and (**L**) CD8 T cells in the spleen. Bar graphs show percent and total number of (**M**) PD-1+CD44+ CD4 and (**N**) PD-1+CD44+ CD8 T cells in the spleen. Data are representative of 3 independent experiments (n = 5 - 8 mice per group). Data are presented as mean ± SEM. Asterisks indicate significant differences (p < 0.05) between ND- and WD-fed mice. (**O**) Representative confocal images of Foxp3+ cells (green) in the liver tissue sections from disease controls and NASH patients (n = 10 subjects per group). Nuclei are stained blue.

### Myeloid cells promote hepatic inflammation and fibrosis in the absence of lymphoid cells

To examine the contribution of adaptive immune cells in promoting NAFLD progression, we fed the WD to *Rag1^-/-^* mice that lack functional T and B cells (**Fig. 3A**). Following sixteen-weeks of WD-feeding, *Rag1^-/-^* mice gained significant body weight as well as liver weight/body weight and visceral fat weight/body weight ratios were also significantly higher in WD-fed *Rag1^-/-^* mice relative to ND-fed mice (**Fig. 3B**). Histologic images of the liver tissue sections stained with H&E revealed micro and macro vesicular steatosis, portal inflammatory infiltrates and ballooning degeneration of hepatocytes similar to WT mice fed the WD for sixteen-weeks (**Fig. 3C** and **Fig. 1F**). Liver injury in WD-fed *Rag1^-/-^* mice were further validated by higher serum AST and ALT levels relative to the controls suggesting that WD-feeding induced significant hepatic injury in the *Rag1^-/-^* mice (**Fig. 3D**). Transcript levels of key inflammatory cytokines, TNFα and IL-1β and chemokine, MCP1 were also significantly higher in the WD-fed *Rag1^-/-^* mice relative to ND-fed controls (**Fig. 3E**). As shown in **Fig. 3F**, WD-fed *Rag1^-/-^* mice also developed extensive pericentral, periportal, and sinusoidal fibrosis. Increased hepatic fibrosis in WD-fed *Rag1^-/-^* mice was corroborated by significant increase in the transcript levels of key markers of hepatic fibrosis, αSMA, TIMP1 and Col1A1 (**Fig. 3G**).

**Figure. 3.**
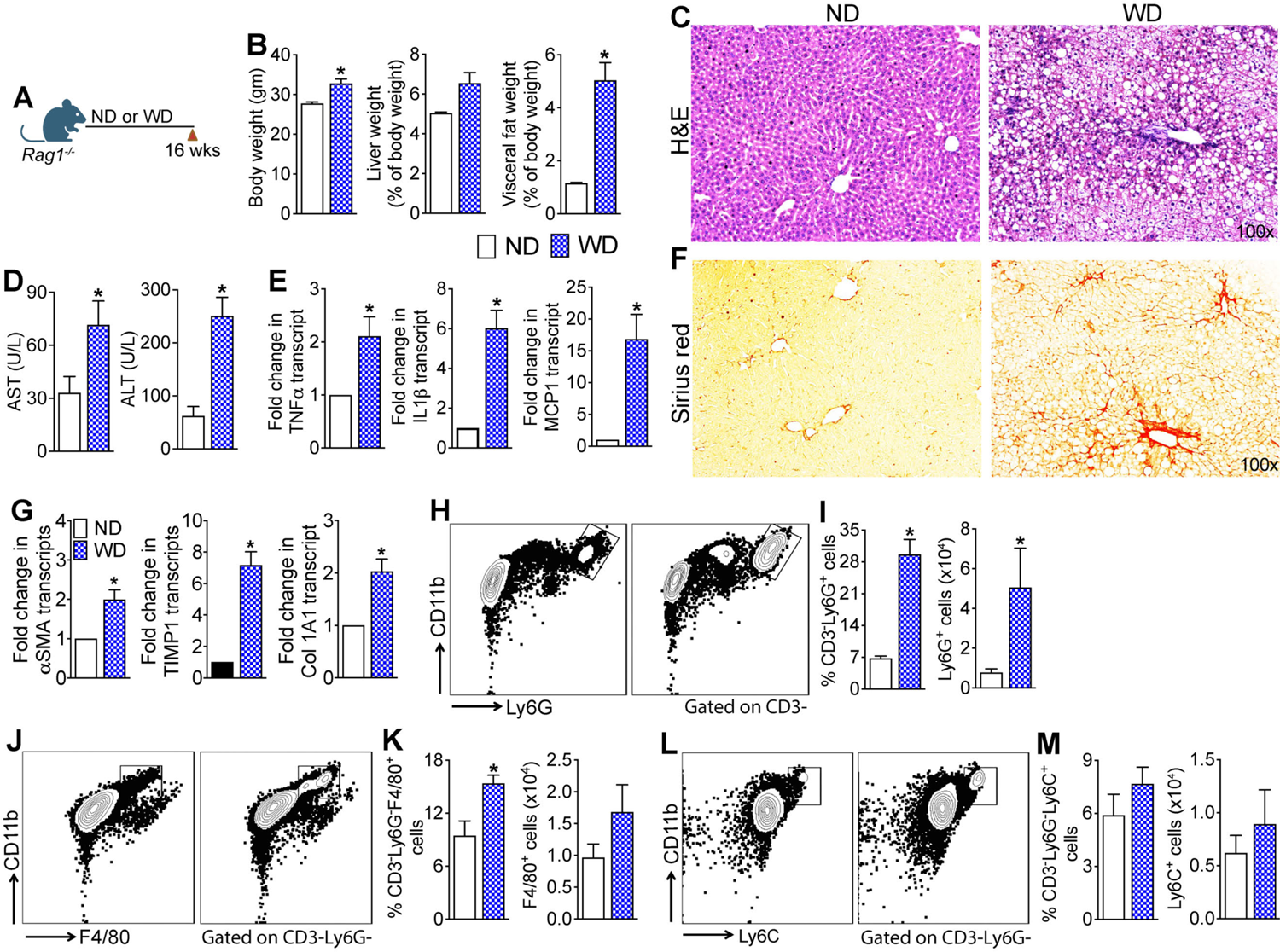
Myeloid cells promote hepatic inflammation and fibrosis in the absence of lymphoid cells. (**A**) Schematic of study design. Rag1 KO mice were fed a ND or WD for sixteen-weeks. (**B**) Body weight and liver and visceral fat weight expressed as percentage of body weight following sixteen-weeks of WD-feeding. (**C**) Representative photomicrographs of Hematoxylin and Eosin (H&E) stained liver tissue sections. (**D**) Serum AST and ALT levels. (**E**) Transcript levels of key inflammatory cytokines in the liver. (**F**) Representative photomicrographs of Sirius Red-stained liver tissue sections. (**G**) Transcript levels of key markers of liver fibrosis. (**H**) Representative flow plots show percent of neutrophils (CD11b+Ly6G+ cells) and (**I**) bar graphs show percent and total numbers of neutrophils in the liver. (**J**) Representative flow plots show percent of macrophages (CD11b+F4/80+ cells) and (**K**) bar graphs show percent and total number of macrophages in the liver. (**L**) Representative flow plots show percent of monocytes (CD11b+Ly6C+ cells) and (**M**) bar graphs show percent and total number of monocytes in the liver. Data are representative of 2 independent experiments (n = 5 - 8 mice per group). Data are presented as mean ± SEM. Asterisks indicate significant differences (p < 0.05) between ND- and WD-fed mice.

To identify the immune cells contributing to hepatic inflammation and fibrosis in the absence of adaptive immune cells we analyzed hepatic immune infiltrates in the WD-fed *Rag1^-/-^* mice (**Fig. 3H-M**). Our phenotypic analyses revealed significant enrichment of neutrophils (CD3-Ly6G+ cells) (**Fig. 3H-I**) and macrophages (CD3-Ly6G-F4/80+ cells) (**Fig. 3J-K**) but not monocytes (CD3-Ly6G-Ly6C+ cells) (**Fig. 3L-M**) in the liver of WD-fed *Rag1^-/-^* mice relative to ND-fed controls. Collectively, these data suggest that in the absence of adaptive immune cells, myeloid cells promote WD-induced hepatic inflammation and fibrosis resulting in the development and progression of NAFLD.

### Depletion of Foxp3+ regulatory T cells reduces hepatic steatosis but exacerbate WD induced hepatic inflammation

Having established that absence of adaptive immune cells does not proffer protection against WD-induced liver injury and that WD consumption results in enrichment of the systemic and hepatic Treg cells, we next investigated whether Foxp3+ Treg cells play a role in NAFLD progression by suppressing aberrant activation of innate and adaptive immune cells. To specifically investigate the role of Treg cells in NAFLD progression, we used *Foxp3*^DTR^ knock-in mice in which Foxp3^+^ Treg cells express human Diphtheria toxin receptor (DTR) and green fluorescence protein (GFP) under the control of Foxp3 transcriptional regulatory element (Foxp3^DTR^) that allows specific depletion of Treg cells in vivo. Insertion of the IRES-DTR-GFP cassette in the *Foxp3* locus does not affect Foxp3 gene expression or Treg cell function^23^. Additionally, DT induced cell death is apoptotic and therefore does not initiate a pro-inflammatory immune response^28–30^, rendering this a robust model to study the effects of WD-induced inflammation in NAFLD pathogenesis. For this study, a cohort of Foxp3^DTR^ mice fed a WD for sixteen-weeks were randomized to receive weekly intraperitoneal injections (50 μg/Kg) of DT or saline for four weeks starting at week twelve^25^ (**Fig. 4A**). We used this standardized regimen as it is effective in depleting Tregs without inducing significant pathology in non-DTR mice.^25^ As shown in **Supp.Fig. 1A-B**, four weeks of DT treatment was effective in significantly reducing intrahepatic Tregs in the WD-fed Foxp3^DTR^ mice. Histologic analysis of liver tissue sections revealed milder steatosis in Treg depleted (TregΔ) mice fed a WD relative to control mice as well as WT mice fed the WD for sixteen weeks, but more severe hepatic inflammation as indicated by increased infiltration of immune cells (**Fig. 4B**). Serum AST and ALT levels as well as the transcript levels of key proinflammatory cytokines and chemokines, TNFα, IL6, IL-1β, IFNγ, MCP1 and markers of monocytes and macrophages, CD68 and F4/80 were significantly higher in the WD-fed TregΔ mice relative to controls suggesting that depletion of Treg cells exacerbated WD-induced hepatic inflammation and injury (**Fig. 4C-E**).

**Figure. 4.**
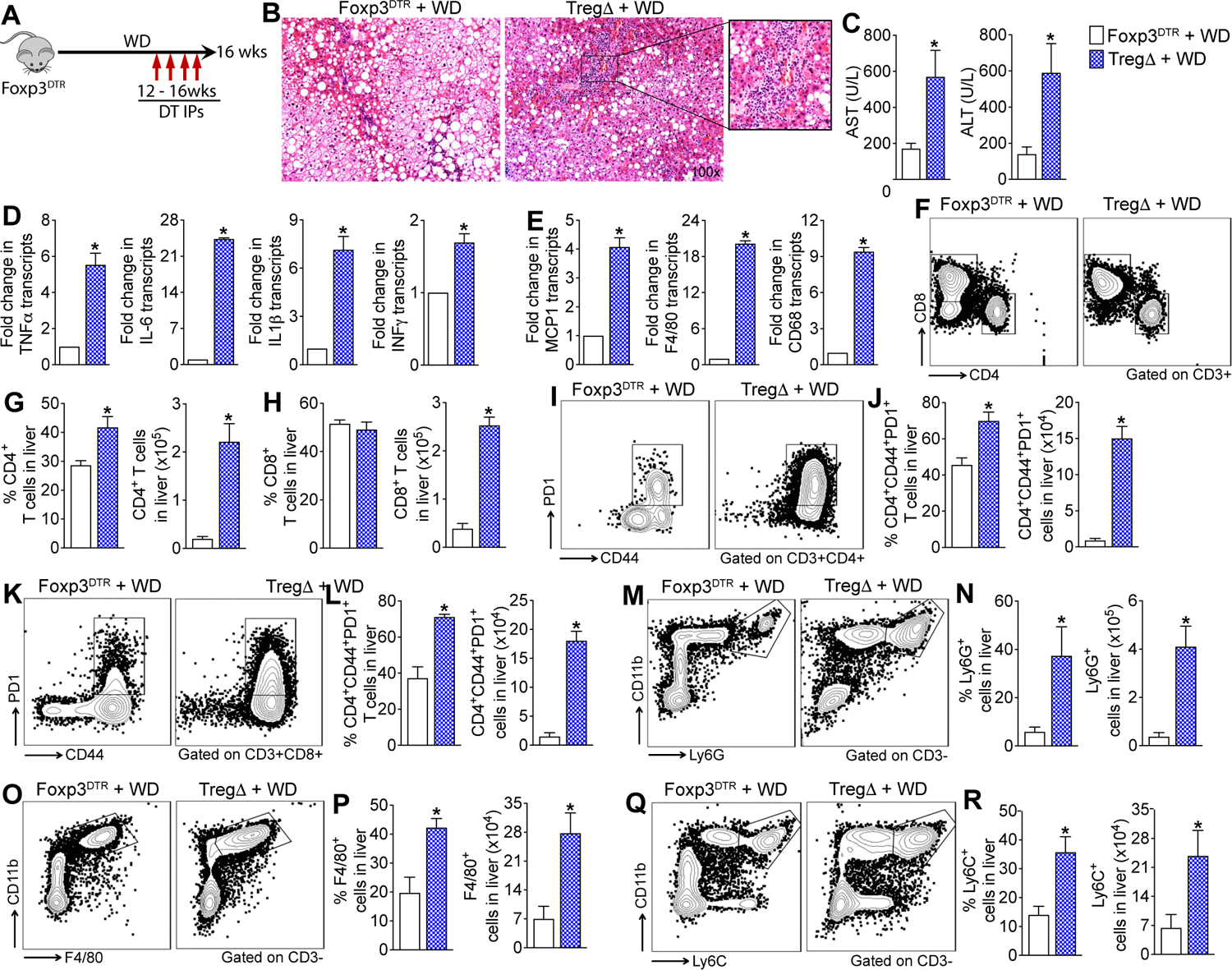
Depletion of Foxp3+ regulatory T cells reduces hepatic steatosis but exacerbates western diet induced hepatic inflammation. (**A**) Schematic of study design. A cohort of Foxp3^DTR^ mice fed a WD for sixteen-weeks were randomized to receive weekly intraperitoneal injections of diphtheria toxin (DT, TregΔ) or saline (controls) for four weeks starting at week twelve. (**B**) Representative photomicrographs of Hematoxylin and Eosin (H&E) stained liver tissue sections. (**C**) Serum AST and ALT levels. (**D-E**) Transcript levels of key inflammatory cytokines, chemokines and markers of monocytes and macrophages in the liver. (**F**) Representative flow plots show percent of intrahepatic CD4 and CD8 T cells. Bar graphs show percent and total number of intrahepatic (**G**) CD4 and (**H**) CD8 T cells. (**I**) Representative flow plots show percent of intrahepatic PD-1+CD44+ CD4 T cells and (**J**) bar graphs show percent and total number of PD-1+CD44+ CD4 T cells. (**K**) Representative flow plots show percent of intrahepatic PD-1+CD44+ CD8 T cells and (**L**) bar graphs show percent and total number of intrahepatic PD-1+CD44+ CD8 T cells. (**M**) Representative flow plots show percent of intrahepatic neutrophils (CD11b+Ly6G+ cells) and (**N**) bar graphs show percent and total number of intrahepatic neutrophils. (**O**) Representative flow plots show percent of intrahepatic macrophages (CD11b+F4/80+ cells) and (**P**) bar graphs show percent and total number of intrahepatic macrophages. (**Q**) Representative flow plots show percent of intrahepatic monocytes (CD11b+Ly6C+ cells) and (**R**) bar graphs show percent and total numbers of intrahepatic monocytes. Data are representative of 3 independent experiments (n = 5 mice per group). Data are presented as mean ± SEM. Asterisks indicate significant differences (p < 0.05) between TregΔ and control mice.

Phenotypic analyses of the hepatic immune infiltrates revealed significant enrichment of intrahepatic CD4 and CD8 T cells in the WD-fed TregΔ mice relative to controls (**Fig. 4F-H**). Further analysis revealed significant enrichment of CD44 and PD-1 double positive activated CD4 and CD8 T cells in the liver (**Fig. 4I-L**). Analysis of the intrahepatic innate immune cells revealed significant enrichment of neutrophils (CD3-Ly6G+ cells), monocytes (CD3-Ly6G-Ly6C+ cells) and macrophages (CD3-Ly6G-F4/80+ cells) in the WD-fed TregΔ mice relative to controls (**Fig. 4M-R**). Collectively, these data demonstrate that Treg suppress aberrant activation and infiltration of adaptive and innate immune cells in the liver of WD-fed mice and protect the liver from WD-induced inflammation and injury.

### Depletion of Foxp3+ regulatory T cells exacerbate WD induced hepatic fibrosis

To determine the effect of Treg depletion on WD-induced hepatic fibrosis, we assessed liver tissue sections for the presence of collagen and quantified transcript levels of key hepatic fibrosis markers (**Fig. 5A-E**). As seen in **Fig. 5B**, WD consumption resulted in more severe hepatic fibrosis in TregΔ mice relative to controls. Consistent with higher hepatic fibrosis in TregΔ mice relative to controls, transcript levels of key liver fibrosis markers, αSMA, TIMP-1 and Col1A1 were also significantly higher in the TregΔ mice (**Fig. 5C-E**). Together, these data demonstrate that Foxp3+ Tregs play a role in mitigation of hepatic inflammation as well as fibrosis in WD-fed mice.

**Figure 5.**
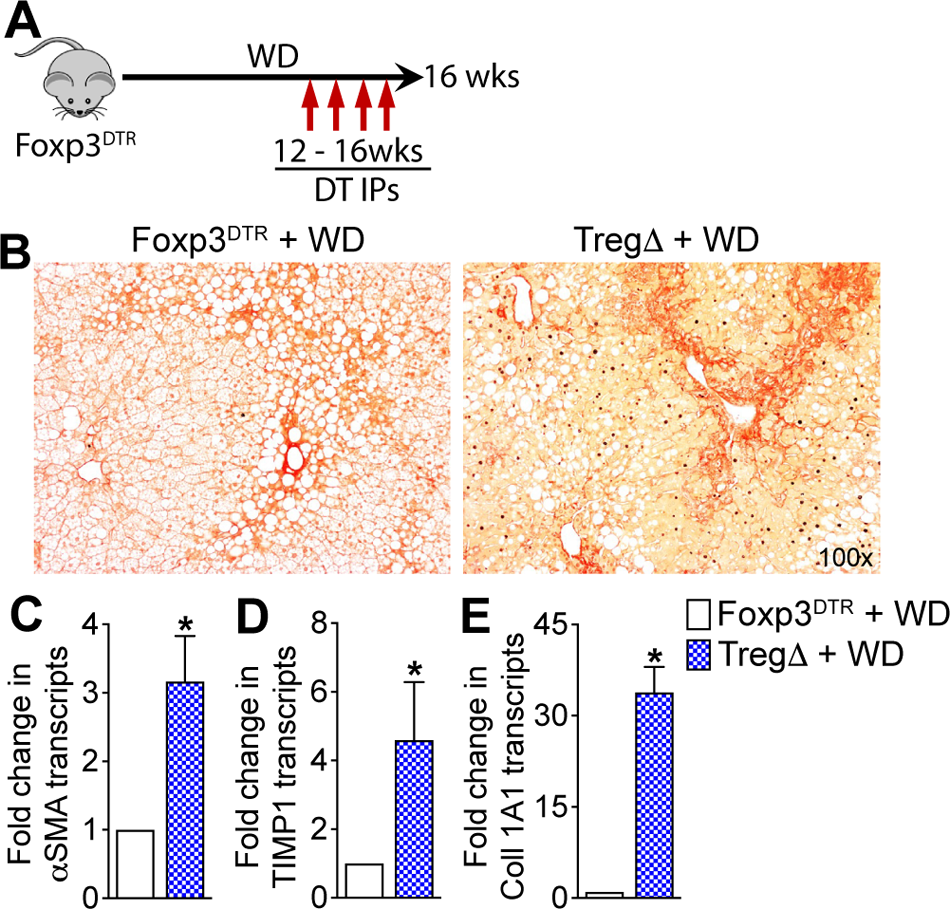
Depletion of Foxp3+ regulatory T cells exacerbate WD induced hepatic fibrosis. **(A)** Schematic of study design. A cohort of Foxp3^DTR^ mice fed a WD for sixteen-weeks were randomized to receive weekly intraperitoneal injections of diphtheria toxin (DT, TregΔ) or saline (controls) for four weeks starting at week twelve. (**B**) Representative photomicrographs of Sirius Red stained liver tissue sections. (**C**) Transcript levels of key liver fibrosis markers in the liver. Data are representative of 3 independent experiments (n = 5 mice per group). Data are presented as mean ± SEM. Asterisks indicate significant differences (p < 0.05) between TregΔ and control mice.

### Induction of Tregs in WD fed mice attenuates hepatic inflammation and steatosis

Having established that Tregs suppress WD-induced hepatic inflammation and fibrosis, we next investigated whether expanding Treg cells will prevent NAFLD progression in WT mice fed a WD for sixteen weeks. We used the established *in vivo* Treg expansion cocktail, recombinant murine IL2 complexed to anti-IL2 mAb (IL2/αIL2 cocktail) to expand Treg cells in mice fed a WD for sixteen-weeks (**Fig. 6A**). As shown in **Fig. 6B-C**, four-weeks of treatment with IL2/αIL2 cocktail resulted in ∼2-fold expansion of intrahepatic Foxp3+ Tregs in the WD-fed WT mice. Assessment of liver histopathology in H&E-stained liver sections revealed marked reduction in hepatosteatosis as well as hepatic inflammation in WD-fed WT mice treated with IL2/αIL2 cocktail relative to saline treated animals (**Fig. 6D**). Reduced hepatic inflammation in the IL2/αIL2 cocktail treated mice was further validated by a significant reduction in the transcript levels of TNFα, MCP-1 and F4/80 in the liver, but no changes in IL-1β was observed (**Fig. 6E-H**). Analysis of the intrahepatic immune cells revealed that the treatment with IL2/αIL2 cocktail significantly increased the percentage of CD4 T cells but did not alter the total number of CD4 T cells in the liver (**Fig. 6I-J**). Interestingly, IL2/αIL2 cocktail treatment significantly reduced both the percentage and total number of intrahepatic CD8 T cells (**Fig. 6K**). Treatment with IL2/αIL2 cocktail also reduced the percentages and total numbers of Ly6C+ monocytes and F4/80+ macrophages, however the reduction in the percentages of monocytes and macrophages did not reach statistical significance (**Fig. 6L-M**). No changes in the percentage and total numbers of Ly6G+ neutrophils were observed between IL2/αIL2 cocktail vs saline treated mice (**Fig. 6N**). Collectively these data suggest that augmenting hepatic Treg population protects mice from WD-induced inflammation and steatosis by reducing the populations of adaptive and innate immune cells in the liver.

**Figure 6.**
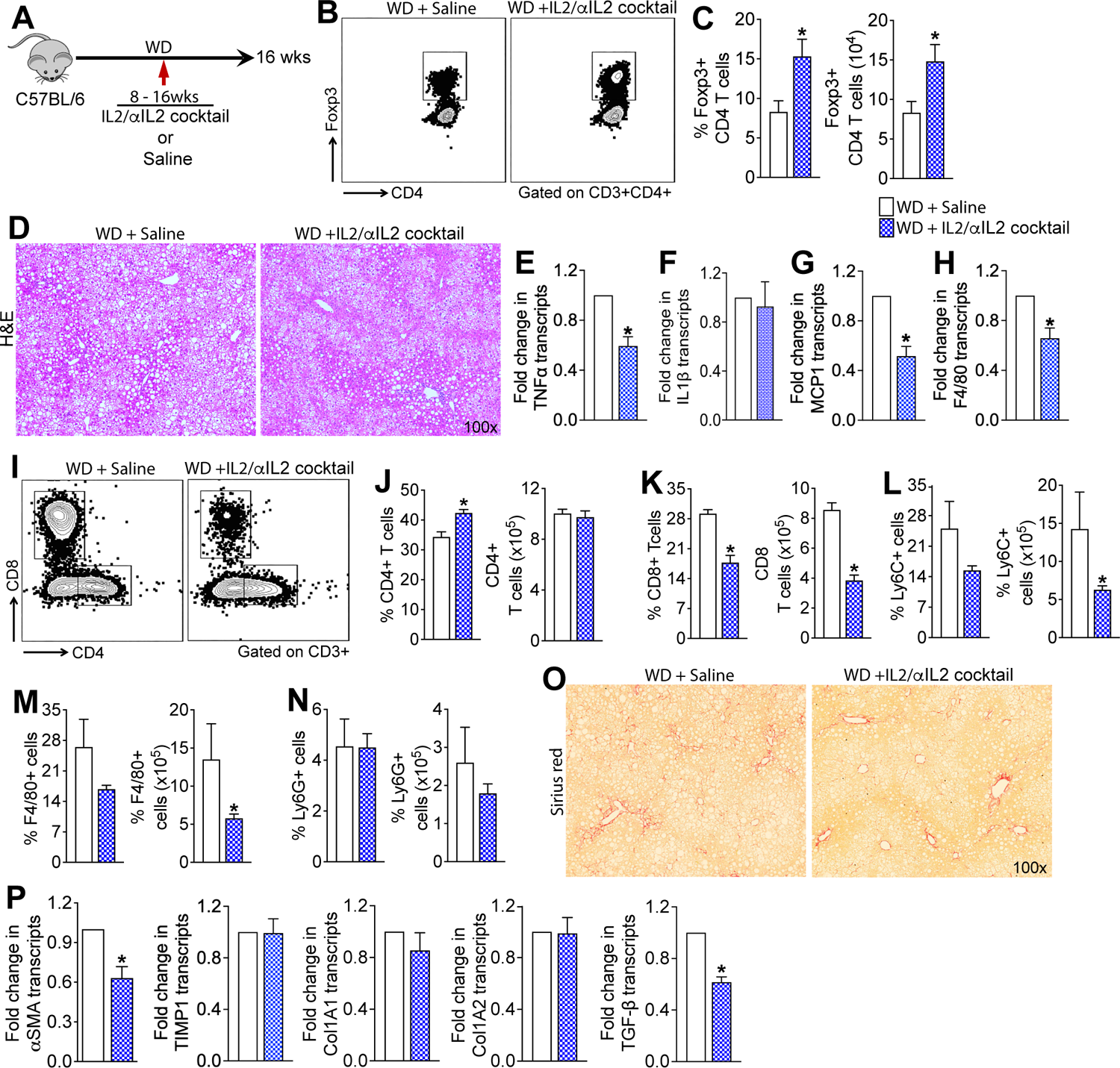
Induction of Tregs in western diet fed mice attenuates hepatic inflammation and fibrosis. (**A**) Schematic of study design. A cohort of WT mice fed a WD for sixteen-weeks were randomized to receive twice weekly injections of Treg induction cocktail consisting of recombinant murine IL2 complexed to anti-IL2 antibody or saline for four weeks starting at week twelve. (**B**) Representative flow plots show percent of intrahepatic Foxp3+ CD4 T cells and (**C**) bar graphs show percent and total numbers of intrahepatic Foxp3+ CD4 T cells. (**D**) Representative photomicrographs of Hematoxylin and Eosin (H&E) stained liver tissue sections. (**E-H**) Transcript levels of key inflammatory cytokines, chemokines and marker of macrophages in the liver. (**I**) Representative flow plots show percent of intrahepatic CD4 and CD8 T cells. Bar graphs show percent and total numbers of intrahepatic (**J**) CD4 and (**K**) CD8 T cells. Bar graphs show percent and total numbers of intrahepatic (**L**) neutrophils (CD11b+Ly6G+ cells), (**M**) monocytes (CD11b+Ly6C+ cells) and (**N**) macrophages (CD11b+F4/80+ cells). (**O**) Representative photomicrographs of Sirius red stained liver tissue sections. (**P**) Transcript levels of key liver fibrosis markers in the liver. Data are representative of 2 independent experiments (n = 5 mice per group). Data are presented as mean ± SEM. Asterisks indicate significant differences (p < 0.05) between saline and IL2/αIL2 cocktail treated mice.

### Treg expansion therapy protects mice from WD-induced liver fibrosis

Assessment of collagen deposition in Sirius red-stained liver sections revealed marked reduction in collagen deposition in the WD-fed WT mice treated with IL2/αIL2 cocktail relative to saline treated animals (**Fig. 6O**). Consistent with lower hepatic fibrosis in IL2/αIL2 cocktail treated animals relative to controls, transcript levels of key liver fibrosis markers, αSMA and TGF-β were significantly lower in the liver, but no differences in TIMP-1, Col1A1 and Col1A2 were observed (**Fig. 6P**). Together, these data demonstrate that augmenting hepatic Treg population protects mice from WD-induced hepatic fibrosis.

### Dietary glucose and palmitate impair hepatic Treg function

Analysis of the intrahepatic Foxp3+ Treg cells revealed that a significantly lower percentage of the Tregs cells in the liver of the WD-fed mice, relative to the ND-fed controls, expressed the proliferation marker Ki67 suggesting that WD-consumption reduces Treg proliferation in the liver (**Fig. 7A-B**). Since PD-1 expressing Treg cells are less proliferative and immunosuppressive^31^, we examined PD-1 expression in the hepatic Tregs cells following WD-feeding. We found that a higher percentage of Foxp3+ Treg cells in the liver of WD fed mice expressed PD-1 relative to the ND-fed mice (**Fig. 7C-D**). These data suggest that WD-feeding impairs hepatic Treg function and provide an explanation for the inability of Tregs to suppress WD-induced hepatic inflammation.

**Figure. 7.**
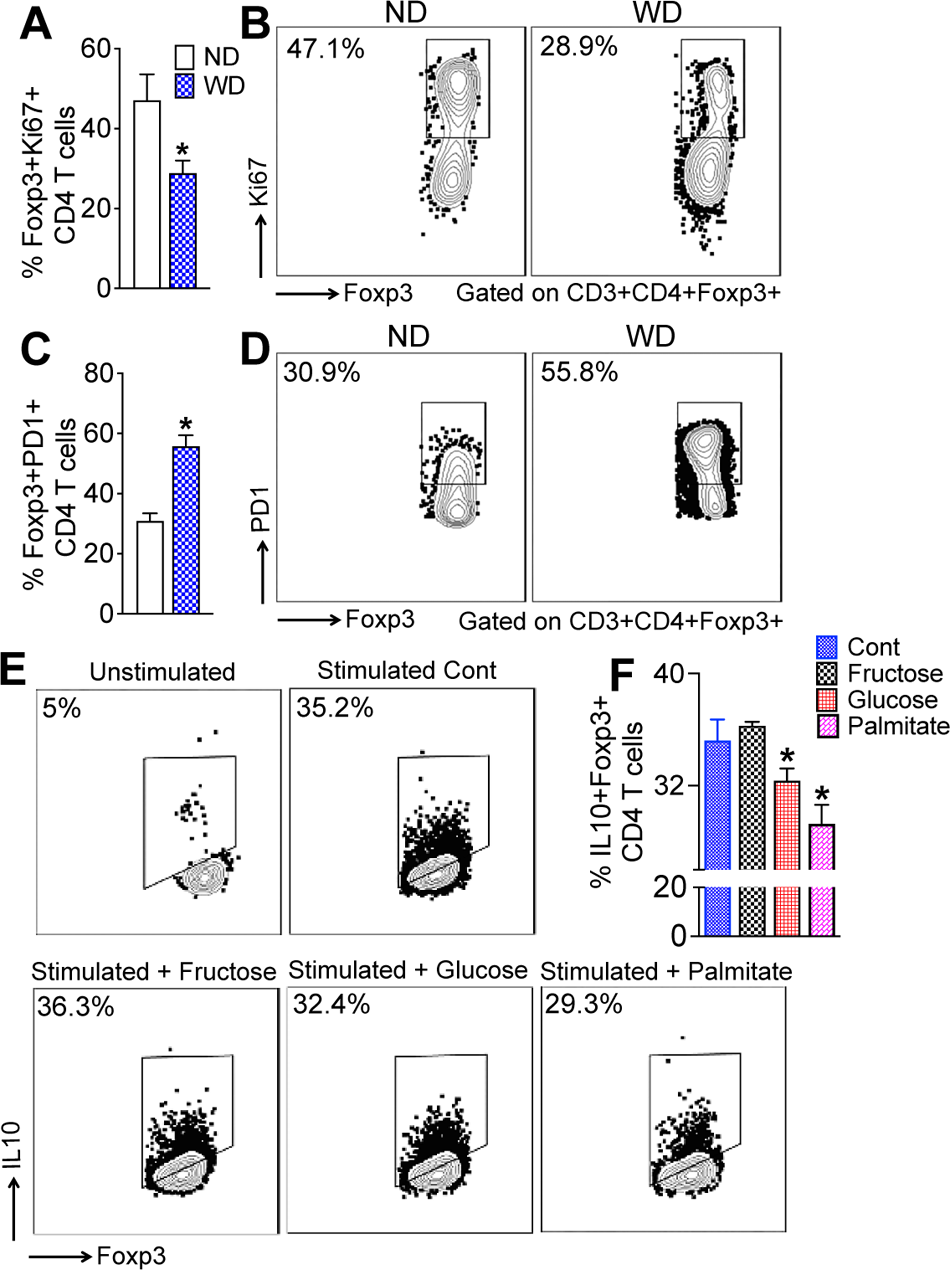
Dietary glucose and palmitate impair hepatic Treg function. (**A**) Bar graphs and (**B**) representative flow plots show percent of FoxP3+ Ki67+ CD4 T cells in the liver of WT mice fed a ND or WD for sixteen-weeks. (**C**) Bar graphs and (**D**) representative flow plots show percent of FoxP3+ PD-1+ CD4 T cells in the liver of WT mice fed a ND or WD for sixteen-weeks. Data are representative of 2 independent experiments (n = 5 mice per group). Data are presented as mean ± SEM. Asterisks indicate significant differences (p < 0.05) between ND and WD fed mice. (**E**) Representative flow plots show percent of FoxP3+ IL10+ CD4 T cells and (**F**) bar graph shows percent of FoxP3+ IL10+ CD4 T cells following treatment with fructose, glucose and palmitate. Splenic CD4 T cells were treated with glucose, fructose or palmitic acid for 24-hours and IL10 production was detected by FACS. Data are representative of 3 independent experiments (n = 3 reps. per treatment). Data are presented as mean ± SEM. Asterisks indicate significant differences (p < 0.05) between stimulated controls and treated cells.

Since a tissue microenvironment that enhances glycolysis and anabolic metabolism in Treg cells impairs proliferative and immunosuppressive ability of Tregs,^32, 33^ we performed ex vivo experiments with mouse primary splenic lymphocytes to determine whether excess glucose, fructose or palmitic acid can impair Treg function (**Fig. 7E-F**). Here, we used IL10 production by Treg cells as a functional read out of immunosuppressive ability of the Treg cells cultured in high concentrations of glucose, fructose or palmitic acid. We found that the percentage of IL10+ Treg cells were significantly reduced when splenic lymphocytes were cultured in high concentration of glucose or palmitic acid (**Fig. 7E-F**). Lymphocytes cultured in high level of fructose did not affect the percentage of IL10+ Tregs (**Fig. 7E-F**). Together, these data suggest that dietary glucose and palmitic acid impairs IL10 production by hepatic Tregs resulting in the inability of Tregs to suppress WD-induced hepatic inflammation in NAFLD.

## DISCUSSION

In this study, we utilized orthogonal approaches of conditional depletion of T cells, targeted Treg diminution, and *in vivo* Treg induction to elucidate the function of Tregs in NAFLD progression. Our data support three major conclusions; first, that Tregs actively infiltrate the liver during progression of NAFLD and suppress activation of innate and adaptive hepatic immune cells thereby resolving inflammation. Second, the immunosuppressive role of Tregs attenuates liver injury preventing rapid progression of NAFLD. Thirdly, however, functional impairment of Tregs in NAFLD, by dietary sugars and fatty acids, results in unfettered immune activation perpetuating chronic inflammation driving disease progression. Our findings therefore resolve an important conundrum in the field of NAFLD about the importance of Tregs in preventing disease progression and suggest that targeted approaches to restore Treg function may prove to be therapeutic in NAFLD.

Our studies using a dietary mouse model of progressive NAFLD show significant accumulation of effector and regulatory T cells in the liver, highlighting their association with hepatic inflammation and fibrosis in NAFLD. Significant increase in the activation of T cells is also observed in the spleen suggesting that WD induces systemic activation of T cells and activated T cells are recruited to the liver. These findings are in agreement with our previous reports demonstrating a role of integrin α_4_β_7_ and its ligand mucosal addressin cell adhesion molecule (MAdCAM)-1 mediated recruitment of effector T cells in the NAFLD liver ^4^. WD consumption also induces systemic increase in Tregs and a larger population of Tregs accumulate in the liver. These findings, along with our previous reports and reports from other groups, support a role of effector T cells and Tregs in NAFLD.

Our first approach to parse out the role of adaptive immune cells in NAFLD was to use Rag-1 KO mice that lack functional T and B cells. Recent reports of both protection and exacerbation of hepatic injury in Rag-1 KO mice led us to reexamine NAFLD development in this model and understand the effect of diet on the myeloid compartment in the absence of functional adaptive immune cells.^19, 34, 35^ We found that Rag-1 KO mice fed a WD for sixteen weeks develop more severe hepatic inflammation and fibrosis compared to WT mice. As expected, hepatic injury in WD-fed Rag-1 KO mice appeared to be driven by the myeloid cells specially neutrophils and macrophages suggesting that the adaptive immune cells protect liver from diet-induced liver injury by suppressing exorbitant activation of myeloid cells. We do not have an explanation for the opposing results in the previous reports. However, a review of the literature revealed that protection from diet-induced liver injury was observed when methionine choline deficient diet (MCD) or choline deficient high fat diet (CD-HFD) was used to induce live injury.^35, 36^ Contrarily, exacerbation of liver injury was observed when WD or high fat diet was used to induce liver injury.^19^ This suggest that the diet influences the immune response resulting in conflicting outcomes observed by different groups.

The overwhelming evidence for immunosuppressive function of Tregs in various inflammatory diseases and its reputation as the major anti-inflammatory immune cells have been challenged by recent reports of a proinflammatory role of Tregs in NAFLD pathogenesis.^19–21^ In contrast to the recent reports of a proinflammatory role of Tregs in NAFLD, our findings demonstrate that targeted Treg depletion exacerbates, while in vivo Treg augmentation reduces WD-induced hepatic inflammation and fibrosis slowing NAFLD progression. These data support an immunosuppressive role for Tregs in NAFLD. In contrast to our findings, a recent study reported protection from CD-HFD-induced hepatic injury in targeted Treg depleted mice and in mice treated with anti-CD25 antibody.^20^ As noted above, the proinflammatory role of Tregs in the targeted Treg depleted mice may have to do with the use of CD-HFD diet. However, it is not surprising that anti-CD25 antibody treatment protected mice from CD-HFD-induced hepatic injury as effector T cells also express CD25 and anti-CD25 antibody also depletes CD25 expressing effector T cells.^22^ For similar reasons, adoptive transfer of CD4+CD25+ T cells may exacerbate hepatic injury.^21^ Furthermore, based on the tissue microenvironment CD25+ CD4 T cells can differentiate into effectors or Tregs and in the absence of in-depth analysis of the fate of the transferred cells it is challenging to make decisive conclusions.

We found that glucose and palmitate impair the immunosuppressive ability of hepatic Tregs, leading to chronic activation of effector cells in NAFLD. Tregs in the NAFLD liver express PD-1 and are less proliferative, suggesting a suppressive liver microenvironment. The compensatory increase in Tregs in NAFLD liver helps regulate immune effector cell activation, but fails to fully resolve hepatic inflammation and fibrosis, leading to disease progression. Our findings, therefore, suggest that harnessing Treg function could be a therapeutic strategy for preventing and treating NAFLD.

