## Supplementary Data for "Western diet dampens T regulatory cell function to fuel hepatic inflammation in nonalcoholic fatty liver disease"

***Contributed equally**

TABLE OF CONTENTS

Supplementary methods ……………………………………………………..……..……2

Supp. Fig. 1 ………………………………………………………………………..4

### SUPPLEMENTARY METHODS

**Histopathology.** Formalin fixed, paraffin embedded liver tissue sections were stained with hematoxylin and eosin (H&E) and Sirius Red as described previously (10). Images were captured using a Zeiss Light Microscope (Zeiss, Jena, Germany).

**Immunofluorescence microscopy.** Formalin fixed, paraffin embedded liver tissue sections were stained for immunofluorescence microscopy as described previously.[1] Sections were visualized using an Axioskop 2 plus laser confocal microscope. Following antibodies were used for human tissue, Foxp3 (cat. No PIPA585236, Thermo Fisher Scientific, Rockford, IL).

**Serological analysis.** Serum alanine aminotransferase (ALT) and aspartate aminotransferase (AST) concentrations were measured using an AST and ALT Activity Assay Kit (Sigma-Aldrich, St. Louis, MO) as described previously.[1]

**Flow cytometric analysis**. Flow cytometric analysis was performed on hepatic lymphocytes as described previously.[1] In short, livers perfused with 1X PBS were digested with 2 mg/mL type IV collagenase (Worthington, NY) to obtain single cell suspension. Lymphocytes were then enriched by Percoll gradient centrifugation, stained with fluorochrome-conjugated antibodies and fixable viability stain (BD Biosciences, San Jose, CA) and acquired on a Cytek Aurora Spectral Cytometer (Cytek Biosciences, Bethesda, MD) equipped with five lasers. The data obtained were analyzed using FlowJo software v X.10 (Tree Star, Inc., Ashland, OR). Fluorochrome-conjugated antibodies against CD3 (clone 17A2), CD4 (clone RM4-5), CD8 (clone 53-6.7), CD44 (clone IM7), and Ly6C (clone AL-21) were purchased from BD Biosciences (San Jose, CA), Foxp3 (clone FJK-16s), Ki67 (clone SolA15), Ly6G (clone RB6-8C5), CD11b (clone M1/70) and Live/Dead Aqua were purchased from Thermo Fisher Scientific (Rockford, IL), and

PD-1 (clone RMP1-30) and F4/80 (clone BM8) were purchased from BioLegend (San Diego, CA).

**Quantitative real-time PCR.** Isolation of total RNA from tissue, synthesizing complementary DNA (cDNA) from RNA and qRT-PCRs were carried out as described previously.[2] Data were normalized to the housekeeping gene 18S rRNA and presented as fold change in gene expression relative to controls.

**REFERENCES.**

1. Rai RP, Liu Y, Iyer SS, et al. Blocking integrin alpha4beta7-mediated CD4 T cell recruitment to the intestine and liver protects mice from western diet-induced non-alcoholic steatohepatitis. J Hepatol 2020.
2. Rahman K, Desai C, Iyer SS, et al. Loss of Junctional Adhesion Molecule A Promotes Severe Steatohepatitis in Mice on a Diet High in Saturated Fat, Fructose, and Cholesterol. Gastroenterology 2016;151:733-746 e12.

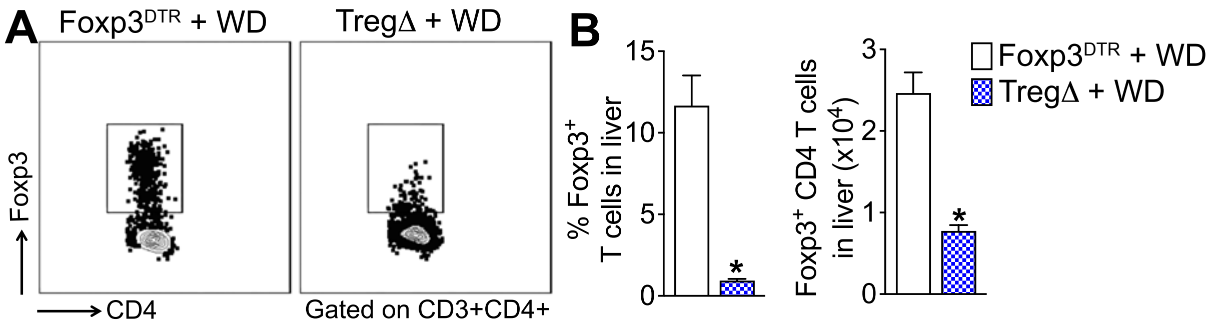


***Supp. Fig. 1. Depletion of Foxp3+ regulatory T cells in Foxp3$^{DTR}$ mice.*** (**A**) Representative flow plots show percent of Foxp3+ regulatory T cells in the liver. Bar graphs show percent and total number of Foxp3+ regulatory T cells in the liver. A cohort of Foxp3$^{DTR}$ mice fed a WD for sixteen-weeks were randomized to receive weekly intraperitoneal injections of diphtheria toxin (DT, TregΔ) or saline (controls) for four weeks starting at week twelve. Data are representative of 3 independent experiments (n = 5 mice per group). Data are presented as mean ± SEM. Asterisks indicate significant differences ($p < 0.05$) between TregΔ and control mice.
